# Guiding Vaccine Efficacy Trial Design During Public Health Emergencies: An interactive web-based decision support tool

**DOI:** 10.1101/252783

**Authors:** Steven E. Bellan, Rosalind M Eggo, Pierre-Stéphane Gsell, Adam J Kucharski, Natalie E Dean, Richard Donohue, Matt Zook, Frank Odhiambo, Ira Longini, Marc Brisson, Barbara E Mahon, Ana Maria Henao-Restrepo

## Abstract

The design and execution of rigorous, fast, and ethical vaccine efficacy trials can be challenging during epidemics of emerging pathogens, such as the 2014-2016 Ebola virus and 2015-2016 Zika virus epidemics. Response to an urgent public health crisis requires accelerated research even as emerging epidemics themselves change rapidly and are inherently less well understood than well-established diseases. As part of the World Health Organization Research and Development Blueprint, we designed a web-based interactive decision support system (InterVax-Tool) to help diverse stakeholders navigate the epidemiological, logistical, and ethical decisions involved in designing a vaccine efficacy trial during a public health emergency. In contrast to existing literature on trial design, InterVax-Tool offers high-level visual and interactive assistance through a set of four decision trees, guiding users through selection of 1) the Primary Endpoint, (2) the Target Population, (3) Randomization, and (4) the Comparator. Guidance is provided on how each of fourteen key considerations–grouped as Epidemiological, Vaccine-related, Infrastructural, or Sociocultural–should be used to inform each decision in the trial design process. The tool is not intended to provide a black box decision framework for identifying an optimal trial design, but rather to facilitate transparent, collaborative and comprehensive discussion of the relevant decisions, while recording the decision process. The tool can also assist capacity building by providing a cross-disciplinary picture of trial design using concepts from epidemiology, study design, vaccinology, biostatistics, mathematical modeling and clinical research ethics. Here, we describe the goals and features of InterVax-Tool as well as its application to the design of a Zika vaccine efficacy trial.

**One Sentence Summary:** An interactive web-based decision support tool was developed to assist in the design of vaccine efficacy trials during emerging outbreaks.

## Introduction

Outbreaks of emerging pathogens pose a major threat to public health and often lead to public health emergencies (PHEs) (*1*). Responding to such outbreaks is extremely challenging because these outbreaks tend to occur as unpredictable events and often accelerate quickly, as evidenced by the recent 2014-2016 Ebola virus (EBOV) and 2015-2016 Zika virus (ZIKV) epidemics. Preparedness activities conducted during inter-epidemic periods are needed to increase the effectiveness and speed of epidemic response activities, including clinical research (*2, 3*). In particular, with the potential for safe and effective vaccines to control or prevent future outbreaks, it is imperative that the international community become better prepared to develop vaccines on rapid time scales (*2*). Even when candidate vaccines are available at the start of an emerging outbreak, as was the case during the 2014-2016 West African EBOV outbreak, their evaluation via phase III efficacy trials must be planned and executed quickly. This requires identification and recruitment of participants at risk of infection; engagement with local communities, and national and international authorities; and consideration of a trial design’s ethicality, feasibility, and acceptability (*4*).

Improving the design of vaccine efficacy trials can be achieved by advance consideration of the epidemiological, logistical, and ethical challenges that might arise in the context of a PHE and how these challenges may interact with vaccine trial design (*5–9*). Challenges may relate to idiosyncrasies of the pathogen, vaccine characteristics, available health systems infrastructure, laboratory capacity, or the sociocultural context of the affected area(s). Decision support tools that help clarify the multidimensional nature of these challenges can play an important role in aiding the trial design process.

The World Health Organization (WHO) Research & Development Blueprint for action to prevent epidemics (WHO R&D Blueprint) aims to integrate research activities into PHE responses, and to improve the design, implementation, and conduct of vaccine evaluation studies in the context of PHEs (*3*). Here, we describe an interactive web-based decision support tool for vaccine trial design, InterVax-Tool (http://vaxeval.com), that was designed as part of the WHO R&D Blueprint working group for vaccine evaluation, that is composed of in infectious disease specialists. The group included content experts such as biostatisticians, trialists, epidemiologists, and mathematical modelers, as well as public health officials from high-, middle‐ and low-income countries. Through an interactive decision tree process, the tool facilitates efficient dialogue amongst diverse decision makers and stakeholders, including ethicists, public health practitioners, policymakers, and national and international authorities.

## InterVax-Tool

### Objectives

While substantial research exists on vaccine efficacy trial design in general (*10*), less guidance exists on trial design for emerging infectious diseases, with much of the available literature developed after the 2014-2016 West African Ebola epidemic (*6, 11–14*). Furthermore, neither scientific literature nor public health agency guidance documents lend themselves well to urgent decision making. This is because the scientific literature is extensive, constantly growing and requires specialist knowledge for interpretation; and because guidance from public health agencies, due to their lengthy and linear nature, obscures the complex interdependencies between decisions (*15*). Many factors–from power estimates, to the logistics of surveillance, to vaccine supply–affect the feasibility of vaccine efficacy trials. Navigating these considerations during emerging outbreaks requires nimble decision making because epidemic dynamics are unpredictable and rapidly changing. This leaves little time to plan around uncertainties or to adapt trial design to the sociocultural context in which an outbreak occurs. This difficult environment requires face-to-face discussion among diverse decision makers, and clear understanding of how decisions on each aspect of trial design affect other downstream choices. Communication between individuals from diverse backgrounds is critical to successful scientific decision making in general (*16*), and this is particularly true during disaster response. Decision support systems have previously been developed to support clinical decision-making (*17*), other aspects of epidemic preparedness, and to guide implementation of epidemic interventions (*18, 19*). However, no decision support system has been developed to assist in the design of phase III vaccine trials during a PHE. The InterVax-Tool was developed to address this gap by satisfying the following objectives:

1. Provide a visual, high-level overview of the key decisions involved in vaccine efficacy trial design during outbreaks constituting a PHE (*4*)
2. Given a particular outbreak context, rapidly highlight the key uncertainties that affect trial design decisions
3. Offer guidance on these decisions from a multidisciplinary perspective
4. Facilitate structured dialogue amongst stakeholders using a common framework
5. Support note-taking and documentation of discussions and decision making
6. Be functional in low bandwidth settings
7. Promote user engagement using principles drawn from usability engineering and user-centered design (*20*)
8. Provide references to literature and other guidance

### Features

InterVax-Tool divides vaccine efficacy trial design into *4 decision topics*: (1) the Primary Endpoint, (2) the Target Population, (3) Randomization, and (4) the Comparator. Major decisions within each of these topics are displayed as a decision hierarchy that allows users to quickly gain a high-level view of the key choices stakeholders must make when designing a vaccine efficacy trial. Users proceed through each of these decision trees one by one, discussing each decision in turn.

InterVax-Tool guides these discussions with content that describes how each decision is affected by *14 key considerations*, which are divided into *4 categories*: (1) pathogen transmission and disease epidemiology (Epidemiology), (2) health systems infrastructure (Infrastructure), (3) vaccine characteristics (Vaccine), (4) and the Sociocultural Context (Sociocultural). The 14 key considerations are listed by category in Figure 1. These factors are expected to be disease‐ and outbreak-specific, comprising an *outbreak scenario*. For the decision under consideration (the “active decision”), InterVax-Tool displays all considerations that are relevant to that decision and how a given consideration may affect this decision. This guidance includes citations to relevant scientific literature as well as links to a more comprehensive WHO R&D Blueprint guidance document (*4*), and allows stakeholders to assess relevant aspects and implications for trial design and to make an informed methodological decision.

**Figure 1.**
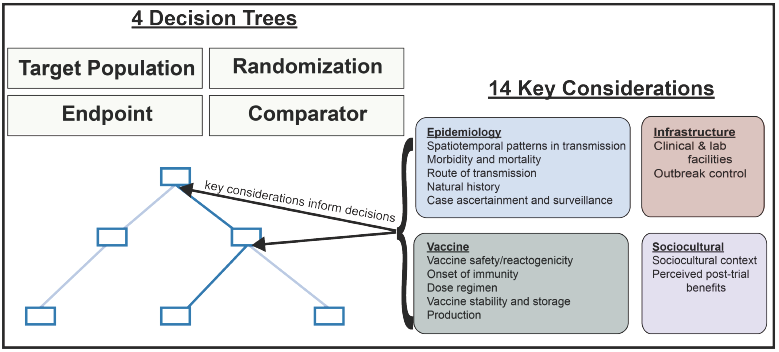
Schematic of InterVax-Tool’s decision process. Within each of 4 decision trees, users navigate a set of hierarchical decisions following guidance on how each of 14 key considerations affect the decision to pick one choice (blue rectangle) over another. During this process users take notes on the scenario under consideration as well as on their justifications for the decisions chosen through the four decision trees.

In addition to guiding decision making with literature-supported content, InterVax-Tool also facilitates continuous discourse by allowing users to take notes both on their decisions taken and on how each key consideration applies to the specific scenario being examined. Users’ notes on their decisions reflect discussion about why a particular decision would be made or, alternatively, the tradeoffs between different choices (Figure 2). Notes entered on key considerations, in contrast, remain consistent across decisions. For instance, if users record notes on a vaccine’s dose regimen while contemplating the primary endpoint, they will see and be able to update these same notes again when considering how dose regimen might impact the choice of comparator arm. This continuity in record-keeping allows users to create a comprehensive record of the epidemic scenario under examination as well as their justification for the best trial design(s). Scenarios and all annotation can be saved to an online database and loaded later, shared it with others, or exported to a static, printable file.

**Figure 2.**
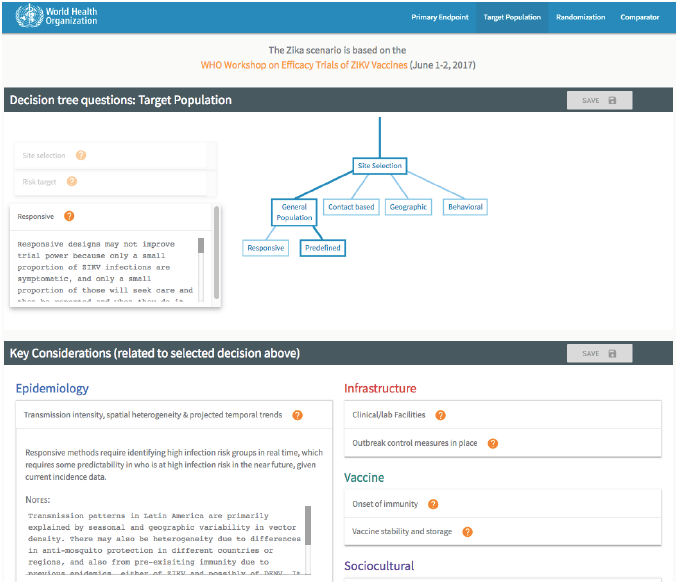
Screenshot of InterVax-Tool at http://vaxeval.com. The decision tree with decisions and decision notes are presented on the top portion of the Tool. The bottom portion provides content on how relevant key considerations impact the active decision in the tree and provides the opportunity for users to take notes on each key consideration, building a description of the scenario during the process.

InterVax-Tool is not intended to give users a single answer on the optimal trial design for a given scenario, but rather to promote organized, efficient, and transparent discussion. In fact, the tool discourages the notion that there may be a single ideal design for any particular scenario and encourages participants to consider tradeoffs between multiple trial design decisions.

A five minute video tutorial gives users a quick introduction to the user interface without requiring that they read a lengthy manual. To allow users to start using the tool as quickly as possible, InterVax-Tool also contains vignettes on Zika virus (ZIKV) and Ebola virus (EBOV) vaccine trial design scenarios. These vignettes include pre-filled notes on both decisions and key considerations so that users can see what they might expect to achieve as a result of using the tool. These vignettes also provide an appropriate starting point when using the tool for capacity building in vaccine trial design.

## Case Study: Zika Vaccine Efficacy Trial Design

InterVax-Tool was piloted at the WHO Workshop on “Efficacy trials of ZIKV Vaccines: endpoints, trial design, site selection” in June 2017 to assist in the design of ZIKV vaccine efficacy trials (*21*). At this meeting, a group of 30 experts used InterVax-Tool to discuss and refine options for potential phase III vaccine trials in Latin America. By using InterVax-Tool, the participants quickly focused their discussion on the three key unknowns in ZIKV vaccine trial planning: 1) ability of assays to distinguish past flavivirus infections, i.e. determining serological positivity for ZIKV separately from dengue virus (DENV; a related flavivirus); 2) forecasted or expected incidence required for feasible sample sizes; and 3) whether to use laboratory-confirmed ZIKV infection (whether asymptomatic or symptomatic) or laboratory-confirmed symptomatic Zika disease as the primary endpoint. The discussion quickly clarified that severe complications from ZIKV infection, e.g. Zika congenital syndrome, or Guillain-Barré syndrome, were very rare outcomes and, therefore, inappropriate as primary endpoints. The ZIKV vaccine pipeline had 45 vaccine candidates at the time of the meeting, so participants agreed to delay focus on vaccine characteristics until candidate vaccines were approaching phase III trials. Meeting attendees agreed that the InterVax-Tool allowed a diverse group of experts to quickly narrow down the range of design possibilities and agree on important knowledge gaps that needed further attention.

### Harmonization with other preparedness activities

In addition to the InterVax-Tool, the WHO R&D Blueprint initiated three other working groups (*22*) to implement complementary initiatives focused on improving vaccine efficacy trial design during emerging outbreaks:

1. A detailed guidance document with expanded descriptions of the key considerations for vaccine study design in general and as applied to WHO R&D Blueprint-designated priority diseases (i.e. diseases evaluated as likely to cause a future PHE) (*4*).
2. A transmission model and trial design simulator capable of addressing a range of diverse questions related to efficacy trials during PHEs (*23*).
3. A set of pre-planned trial generic design protocols from which investigators can build for future trials evaluating vaccines to prevent Blueprint priority diseases.

There is harmonization between each of these initiatives, although each can stand alone. The InterVax-Tool decision support tool refers extensively to the guidance document (*4*), and unresolved questions that arise from InterVax-Tool may be addressable with the trial simulator tool in the second initiative. To maximize the utility of the InterVax-Tool, feedback was iteratively solicited from the other three working groups to maximize the coherence of the entire set of products. For instance, the hierarchical organization of trial design into the four decision trees described above was achieved through iterative design and feedback during face-to-face meetings of all four working groups. The InterVax-Tool, guidance document, and generic protocols all reflect the final consensus hierarchy.

## Limitations

The guidance presented within InterVax-Tool reflects the collective expertise of this working group and its attempt to summarize the scientific literature on vaccine efficacy trial design. We encourage user feedback so that content can be updated to reflect the state-of-the-art on all facets of trial design.

Because InterVax-Tool is organized in a hierarchical decision tree framework, decisions are reflected as discrete choices between two or more options. In reality, some decisions may fall along a spectrum and not all categories are mutually exclusive. Nonetheless, the categorization scheme used encourages discourse and planning, though it may at times simplify complex aspects of trial design.

Further, because InterVax-Tool aims for simplicity to avoid user fatigue, some trial design options were excluded (e.g., factorial design and interim analysis options). We provide ample reference to the scientific literature on these topics as well as to the complementary, comprehensive WHO guidance document (*4*).

## Conclusion

We created a web-based interactive decision tree tool, InterVax-Tool, to support decision making on vaccine trial design. By focusing users on the key decisions, trade-offs, and interdependencies, InterVax-Tool allows multidisciplinary users to quickly identify the key issues for each outbreak scenario and thereby assist the vaccine response to public health emergencies. This tool was piloted during a consultation to plan a ZIKV vaccine efficacy trial, and the WHO R&D Blueprint plans to use the tool in preparedness exercises to plan trials for Blueprint priority diseases. An important feature of the tool is the ability to transmit annotated decision trees and design justifications between users. For example, investigators could use the tool to provide their justification for a particular trial design to regional public health authorities. The tool may be similarly useful to vaccine manufacturers thinking about trial scenarios for a vaccine early in the pipeline. Finally, the tool may be useful in capacity building, providing students of trial design and emerging infections with a lens into this complex decision process. The conceptual design of this tool may be applicable to many other aspects of public health or other disciplines in which rapid, transparent, highly technical, and interdisciplinary decision making is necessary to address an urgent problem.

## Acknowledgements

We gratefully acknowledge input from Peter Dull, John Edmunds, Peter Smith, and Laura Rodrigues in addition to the rest of the WHO R&D Blueprint Working Group on Vaccine Study Design and the attendees of the WHO Consultation on Zika Vaccine Trial Design who piloted the tool.

## Author Contributions

SEB conceived the idea of a vaccine efficacy trial design decision tree and wrote the first draft of the manuscript. SEB, RME, PG, and AJK devised the interactive nature of the decision support tool, the hierarchical structure of the decision trees, and developed the guidance content and organization of the tool. AMHR convened the WHO Blueprint working groups for vaccine evaluation in four occasions since March 2016. AMHR, FO, MB, BEM, IL and ND contributed to the guidance content and structure. All authors contributed to the iterative development of the tool, reviewed the guidance content within the tool, and reviewed the final manuscript.

